# Pupillary responses to affective words in bilinguals’ first versus second language

**DOI:** 10.1101/506998

**Authors:** Wilhelmiina Toivo, Christoph Scheepers

**Affiliations:** Institute of Neuroscience and Psychology, University of Glasgow, Glasgow, UK

**Keywords:** Bilingualism, word processing, emotion, pupillometry

## Abstract

Late bilinguals often report less emotional involvement in their second language, a phenomenon called *reduced emotional resonance in L2*. The present study measured pupil dilation in response to high- versus low-arousing words (e.g., *riot* vs. *swamp*) in German-English and Finnish-English late bilinguals, both in their first and in their second language. A third sample of English monolingual speakers (tested only in English) served as a control group. To improve on previous research, we controlled for lexical confounds such as length, frequency, emotional valence, and abstractness – both within and across languages. Results showed no appreciable differences in post-trial word recognition judgements (98% recognition on average), but reliably stronger pupillary effects of the arousal manipulation when stimuli were presented in participants’ first rather than second language. This supports the notion of reduced emotional resonance in L2. Our findings are unlikely to be due to differences in stimulus-specific control variables or to potential word-recognition difficulties in participants’ second language. Linguistic relatedness between first and second language (German-English vs. Finnish-English) was also not found to have a modulating influence.

## Introduction

It is estimated that more than half of the human world population are *bilinguals*, i.e. people who regularly use more than one language in their daily lives. Definitions of bilingualism actually vary across the field. According to a more narrow view, only people who have learnt their second language (L2) from an early age, and thus achieved a native-like fluency in L2, are classified as bilinguals (for discussion, see (1)). A wider definition of bilingualism focuses more on linguistic exposure and active language use, characterising bilinguals as people who regularly use two languages in everyday settings, regardless of the level of fluency they achieve in L2 (1). In line with many other studies in the psycholinguistic literature, the present paper adopts the latter, wider definition of bilingualism. We can further distinguish between two broad subtypes of bilinguals, namely ‘late’ versus ‘early’ bilinguals, by making reference to age of acquisition of L2: A *late bilingual* is a person who did not acquire their second language from birth and therefore has been initially exposed to L2 at later stages of linguistic development, usually through classroom instruction and/or immigration. The term *early bilingual*, by contrast, describes a person who has been exposed to L2 from a very early age, often in the context of parents from different linguistic backgrounds (1). The present work will primarily focus on the first, *late bilingual* subtype.

Late bilinguals often report ‘feeling less’ in their second language, implying that the language acquired later in life does not bear the same emotional weight as the language one initially grew up with (e.g., (2)). As a consequence, late bilinguals tend to prefer using their first language when expressing strong emotions (e.g.,(2, 3)), while on the other hand finding it easier to talk about ‘awkward’ topics (such as sexuality or situations in which they felt embarrassed) in their L2 (4). These findings can be characterised by a feeling of *emotional distance* in L2: Despite the fact that bilinguals may be highly proficient in their second language, the ‘feeling behind the words’ is often reported to be missing in L2, potentially making it easier to communicate in L2 when emotional investment is costly or less desirable (5). This phenomenon is generally referred to as *reduced emotional resonance in L2*.

Reduced emotional resonance in L2 appears to have important implications for decision making and other types of behaviour. For example, Puntoni and colleagues (6) found that bilinguals rated advertisement slogans presented in their L1 as more emotional in comparison to adverts presented in their L2. Keysar and colleagues (7) used a modified version of the *Asian Disease Problem* (8) in which participants have to choose between alternative intervention programmes when faced with a hypothetical disease-outbreak scenario. Participants are typically prone to framing effects in this task, i.e. they tend to be less risk-averse when the problem solution is framed in terms of *number of lives lost* as opposed to the complementary *number of lives saved*. Across three experiments, Keysar and colleagues (7) found that such framing effects disappeared when the problem was presented in participants’ L2, and that bilinguals applied more systematic thinking in L2 compared to when the problem was presented in their L1. More recently, Costa and colleagues (9) were able to replicate these findings, and more interestingly, they showed (via comparison of problem types with different degrees of emotional involvement) that reduced emotional resonance in L2 could indeed play an important role in explaining the more rational decision making in L2-framed problems.

More direct attempts at measuring reduced emotional resonance in bilinguals’ L2 actually prove to be quite challenging, due to the fact that comprehension and production of L2 generally requires more cognitive effort than that of L1. For example, semantic processing is typically slower and less automatic in L2 than in L1 (e.g., (10)) and the speed/accuracy of word processing in L2 is often not as good as in L1 (e.g. (11)). This may create potential confounds when measuring reduced emotional resonance in L2, bearing the risk that such effects may be due to cognitive effort instead of reduced emotional resonance per se. The use of cognitive tasks, or tasks based on implicit processing of linguistic stimuli, appears particularly prone to this type of confound. Indeed, findings based on such tasks have remained somewhat inconclusive in terms of reduced emotional resonance in L2. For example, Segalowitz and colleagues (10) found smaller effects of emotional valence in L2 as opposed to L1 in an implicit association task, which was interpreted as evidence for differences in emotional processing between L1 and L2. More recent studies utilising paradigms such as the Emotional Stroop task (12–14) have found larger interference effects in L1, suggesting that word processing is more affected by emotion in L1 as opposed to L2. However, other studies based on cognitive paradigms have failed to detect any evidence for reduced emotional resonance in L2. For example, Kazanas and Altarriba (15) found no differences between L1 and L2 in an affective priming lexical decision task. Likewise, Eilola, Havelka and Sharma (16), Sutton and colleagues (17), as well as Eilola and Havelka (18) all found identical interference effects between L1 and L2 in an Emotional Stroop task, i.e. there was no clear indication of reduced emotional resonance in L2.

Another route to finding evidence for reduced emotional resonance in L2 has been to use physiological measurements of affect. Compared to survey-type studies or tasks based on cognitive processing of linguistic material, physiological measures offer a more direct and objective way of detecting reduced emotional resonance in L2. This is because such measures reflect spontaneous responses to emotional content without relying on metalinguistic judgement or addressing emotion ‘indirectly’ via its mediating effect on cognitive effort. Indeed, pioneering work based on skin-conductance (e.g., (19–21)) has provided the perhaps most compelling evidence in support of reduced emotional resonance in L2 to date. Testing both late and early bilinguals, and using both auditory and written stimuli, Caldwell-Harris and colleagues have measured Spanish-English, Turkish-English and Mandarin-English bilinguals’ skin-conductance responses (SCR) to different types of emotionally arousing stimuli. These included taboo phrases like “*She’s a slut”*, as well as insults (e.g. “*You’re so* fat”), childhood reprimands (e.g. “*That’s not nice”)* and endearments (e.g. “I *love you*”). Some of the experiments also included aversive versus positive words. Over several experiments, these emotional stimuli were found to evoke consistently stronger SCRs than neutral control stimuli, but only in participants’ L1 and much less so in their L2 (21). Moreover, the difference in emotional SCR between participants’ L1 and L2 was generally more pronounced for late bilinguals than for early bilinguals (19, 21).

While the above studies generally support the notion of reduced emotional resonance in L2, their findings varied to some extent dependent on type of stimulus and participant population. For example, Caldwell-Harris and colleagues (19–21) found that in late Spanish-English and Turkish-English bilinguals, endearments and taboo words elicited stronger emotional SCRs in L2 than in L1. On the other hand, Mandarin-English bilinguals showed no significant differences in emotional SCRs between languages, except for L2 endearments, which again elicited a higher emotional response than their L1 counterparts. The authors conjectured that these unexpected aspects of the findings could be explained by cross-cultural differences in norms of emotional expression (22).

A potential issue in the above SCR studies (as well as in some of the cognitive studies discussed earlier) is that the stimuli were not very tightly controlled in terms of lexical variables (length, frequency, abstractness, etc.), or in terms of syntactic/pragmatic complexity when multi-word phrases were used. This, again, introduces a number of potential confounds (both within and across languages), making it difficult to separate effects of emotional resonance from those related to cognitive effort in word processing. In the present study, we aimed at eliminating such potential confounds by matching our stimuli on a number of lexical variables, and by avoiding the use of translation equivalents with different lexical characteristics across languages.

The present study will evaluate pupillometry as an alternative indicator of reduced emotional resonance in L2. Pupillometry is a non-intrusive way of examining automatic physiological responses to stimuli (23). Just like skin conductance, pupillary responses are an indicator of activity in the Autonomic Nervous System (ANS), which implies that such responses are not under voluntary control. Pupil dilation and constriction are controlled by the dilator and sphincter muscles, which are mediated by both the sympathetic and parasympathetic divisions of the ANS (24, 25). There is a wealth of research demonstrating that pupillary responses are sensitive to cognitive load associated with various tasks and stimulus characteristics. Examples include attention and memory load (e.g.,(26), and (27, 28) as cited in (29)), processing load associated with simultaneous interpretation (e.g., (29)), and ‘garden-path’ effects in syntactic processing (e.g., (30, 31)), to name but a few. One noteworthy study found that pupillary responses are also sensitive to word retrieval effort in L1 and L2 (11), which will be highly relevant for the present investigations (see further below).

Apart from cognitive load, pupil dilation is also known to be sensitive to affect, and more specifically, to the processing of emotionally arousing stimuli. For instance, Partala and Surakka (25) played highly emotional sound clips to their participants, which were of either negative (e.g., a couple fighting) or positive (e.g., laughter) emotional valence. They found that participants’ pupils dilated more in response to such stimuli than in response to emotionally ‘neutral’ sound clips (e.g., office background noise). Interestingly, emotional valence (positive vs. negative) did not appear to play a major role in the observed pupillary response patterns, leading the authors to conclude that pupil size changes in response to affective stimuli are more strongly driven by emotional arousal (how “exciting vs. calming” the items are to perceivers) than by emotional valence (how “pleasant vs. unpleasant” the items are to perceivers). Bradley and colleagues (24) found very similar pupillary response patterns using emotional pictures instead of sound clips.

A more recent, and highly relevant example is a study by Iacozza, Costa, and Duñabeitia (32). They presented two types of sentences to Spanish-English bilinguals. The first type of sentence contained high-arousing negative words (e.g., *hostile terrorist*) in various critical positions throughout the sentence, whereas the second contained low-arousing words of neutral valence (e.g., *civil receptionist*) in the same critical positions. The sentences were either in Spanish or in English, such that half of the Spanish-English bilinguals were reading sentences materials in L1 whereas the other half were reading sentence materials in L2 (randomised between-subjects design). During reading, participants’ pupillary responses were continuously monitored, and at the end of each trial, participants had to provide emotionality ratings on a 7-point Likert scale. The pupillometry data (obtained during on-line reading) showed evidence for reduced emotional resonance in L2: Sentences with high-arousing, negative content evoked reliably more dilated pupils than sentences with low-arousing, neutral content, but this effect was significantly reduced for participants who read the materials in English (L2). Interestingly, the offline ratings did not reveal such an interaction between sentence type and target language: Sentences with high-arousing, negative content were rated as more emotional than sentences with low-arousing, neutral content, but this difference was the same between the two target language conditions (L1 vs. L2). These findings support the notion of reduced emotional resonance in L2 and further suggest that ‘direct’ physiological measurements such as pupillometry are more sensitive to variation in emotional resonance than measures based on explicit metacognitive judgement.

The present pupillometry study builds upon previous work on reduced emotional resonance in L2, with the aim to implement some methodological improvements as well as addressing further theoretical questions.

Instead of high- vs. low-arousal words embedded in sentences (32), we used isolated words as target stimuli in our study. Each word was centrally presented for a maximum of 250 ms before being replaced with a visual mask for 1.7 seconds after word-offset. This had the advantage of being able to minimise the impact of eye-movements on the measurement of pupil size (note that dependent on eye-tracker setup, obtained pupil-sizes usually vary as a function of eye-position). Second, we used a *within-subjects* design to address reduced emotional resonance in L2: Our bilingual participants were tested in both L1 and L2 over two separate sessions (order counterbalanced) such that each participant served as their own control when assessing the effect of target language on affective pupillary responses; a sample of monolingual native speakers of the bilinguals’ L2 served as an additional control group to ensure that the L2 stimuli evoked the expected affective response pattern when presented to L1 speakers of that language. Third, we examined whether similarities between L1 and L2 play a role when measuring reduced emotional resonance in L2. Specifically, we tested two groups of late bilinguals, namely native German and native Finnish speakers, both highly proficient in English as their second language. From a comparative linguistics point of view, German and English belong to the same *Germanic* language family; as such, the two languages share some basic vocabulary, including a sizeable number of cognates (words such as *GARDEN* / *GARTEN*, which have similar spelling and virtually the same meaning in both languages) as well as a number of ‘false friends’ (words such as *BALD*, which are spelled the same but mean different things in German and English). Finnish, on the other hand, belongs to the *Finnic-Uralic* language family (sharing properties with Estonian and Hungarian) and is thus rather different from English in terms of basic vocabulary, phonology, morphology, etc. Comparing the two groups of bilinguals therefore allowed us to establish whether ‘linguistic distance’ between L1 and L2 matters for emotional resonance in L2. Fourth, we aimed at controlling for a wider range of potential lexical and semantic confounds in the linguistic materials. Given that pupillary responses were previously found to be sensitive to word retrieval effort in L1 and L2 (11), we matched our stimuli as closely as possible (across languages as well as conditions) in terms of length, lexical frequency and abstractness, using available word-norm databases for each language. In addition, these variables were also included as covariates in our analyses. Finally, participants in the present study were also asked to provide explicit word recognition judgements at the end of each trial, making it possible to determine whether bilinguals were experiencing any problems in accessing the meanings of the words presented in L2.

Based on the findings discussed earlier, we expected that words associated with high emotional arousal would evoke reliably more dilated pupils than words associated with low emotional arousal. Assuming reduced emotional resonance in L2, this affective pupillary response should primarily become manifest when participants are tested in their L1 (i.e., English speakers tested in English, Finnish speakers tested in Finnish, and German speakers tested in German) but not – or reliably less so – when tested in their L2 (Finnish or German bilinguals tested in English). As a reflection of greater cognitive effort associated with second-language processing, we also expected generally more dilated pupils for bilinguals tested in L2 rather than L1. Finally, under the assumption that linguistic distance could modulate reduced emotional resonance in L2, we hypothesised that when tested in L2 (English), German bilinguals might display less of a reduction in the critical pupillary response contrast (high- vs. low-arousal words) than Finnish bilinguals.

## Method

### Participants

One-hundred-two participants (76 females) were recruited from the University of Glasgow student community. Participants were aged between 17 and 33 years (mean age: 22 years) and were awarded with either course credits or £5 subject payment.

Thirty-two participants were monolingual native English speakers who reported English to be the only language they were using on a day-to-day basis. Seventy participants were bilingual native Finnish or native German speakers, respectively, all of whom reported to be highly proficient in English and to use English regularly in everyday communication. Data from six bilingual participants were excluded from the sample, leaving a total of 96 participants for analysis (32 English monolinguals, 32 Finnish-English bilinguals and 32 German-English bilinguals). One of the excluded participants reported having dyslexia; four bilingual participants were excluded because they reported having been exposed to English from early childhood, and that they were as fluent in English as they were in Finnish or German (only *late bilinguals* were included in the final sample); finally, one bilingual participant was excluded because they reported having misunderstood the task.

Across the remaining 64 bilingual speakers, the reported cumulative length of stay in an English-speaking country ranged from three months to about nine years, with an average reported length of stay of about 28 months (Mean ± *SD* for the Finnish-English bilinguals: 30 ± 16 months; for the German-English bilinguals: 27 ± 24 months). The 32 native Finnish bilinguals reported to have started learning English at an age of 8.4 years on average, and for the 32 native German bilinguals, the average reported age at which they started to learn English was 9.1 years.

### Ethics Statement

The research was approved by the University of Glasgow College of Science and Engineering Ethics Committee. Participants gave written informed consent prior to taking part.

### Design and Materials

A shortened version of the Language History Questionnaire (LHQ 2.0; (33)) was administered to bilingual participants to identify the variables reported in the Participant section.

For the stimulus materials of the main experiment, initial candidate words were selected from six different databases. English words were selected from ANEW (34) and the more recent and extensive Warriner, Kuperman, and Brysbeart (35) database. The Finnish materials were selected from two existing databases on arousal and valence in Finnish: the 210-word corpus by Eilola and Havelka (36), and the 420-word corpus by Söderholm, Häyry, Laine, and Karrasch (37). For the German stimuli, BAWL-R (38) and Leipzig Affective Ratings (39) were used to select candidate words. To normalize affective ratings across different databases, scores were transformed into scale-range proportions such that 0 corresponded to the lowest and 1 to the highest point on the arousal scale in any given database. On the normalized scale, words scoring lower than 0.33 were classified as Low Arousal and words scoring higher than 0.66 as High Arousal, respectively. This resulted in an initial pool of 139 candidate words in English (74 High- and 65 Low Arousal), 115 candidate words in Finnish (64 High- and 51 Low Arousal), and 141 candidate words in German (69 High- and 71 Low Arousal).

For each of these initial word candidates, we also recorded its length (in number of characters and number of syllables), emotional valence (on a normalized scale, as above), abstractness (normalized scale), and lexical token frequency (log_10_ per million word counts). Emotional valence ratings were available from the previous corpora. Abstractness ratings for the English words were obtained from the Brysbaert, Warriner and Kuperman (40) database. For the German and Finnish stimuli, abstractness ratings were mostly available from the previously quoted databases. However, some words taken from the Söderholm et al. (37) corpus did not include abstractness ratings in Finnish. In those instances (ca. 10% of the Finnish candidate items), abstractness ratings were estimated via averaging across normalised scores for translation-equivalent words in the English and German corpora. Lexical token frequencies per million were obtained from the *British National Corpus* (English), from *CSC Kielitaajuussanasto* (Finnish), and from *Leipzig Affective Ratings* respectively *Kernkorpus 21* (German).

From the initial set of candidate words, we selected 30 for each language and emotional arousal category such that differences in the control variables (length in characters and syllables, emotional valence, abstractness, and lexical token frequency) were kept reasonably small both across arousal conditions and across languages. Given that bilingual participants were tested twice (once in L1 and in once in L2), we also avoided using direct translation equivalents across the three material sets. Appendix S1 shows the selected items per language and Table 1 summarizes their average item characteristics.

**Table 1.** Item Characteristics. Item characteristics (Means ± *SDs*) of the 30 High- and 30 Low Arousal words selected for each language. Arousal, valence, and abstractness scores are on normalized scales ranging from 0 to 1 (lowest to highest scale point). Lexical frequency is given in log10 per million word counts. Bottom three sections show Anderson-Rubin factor scores of the three principal components extracted from the control variables (letters, syllables, frequency, valence, and abstractness), as explained in the text and in Appendix S2.

|  |  | Language |  |  |
| --- | --- | --- | --- | --- |
|  |  | English | Finnish | German |
| Arousal | High Arousal | $0.74 \pm 0.04$ | $0.70 \pm 0.05$ | $0.75 \pm 0.05$ |
| | Low Arousal | $0.21 \pm 0.05$ | $0.12 \pm 0.03$ | $0.13 \pm 0.03$ |
| Letters | High Arousal | $6.57 \pm 2.05$ | $6.80 \pm 1.69$ | $6.20 \pm 1.63$ |
| | Low Arousal | $5.43 \pm 1.57$ | $5.43 \pm 1.38$ | $5.67 \pm 0.92$ |
| Syllables | High Arousal | $2.27 \pm 0.79$ | $2.77 \pm 0.63$ | $2.03 \pm 0.62$ |
| | Low Arousal | $1.77 \pm 0.63$ | $2.30 \pm 0.54$ | $1.87 \pm 0.43$ |
| Frequency | High Arousal | $1.00 \pm 0.72$ | $0.96 \pm 0.49$ | $1.18 \pm 0.52$ |
| | Low Arousal | $1.36 \pm 0.67$ | $1.30 \pm 0.49$ | $1.44 \pm 0.55$ |
| Valence | High Arousal | $0.40 \pm 0.28$ | $0.34 \pm 0.34$ | $0.32 \pm 0.17$ |
| | Low Arousal | $0.56 \pm 0.11$ | $0.51 \pm 0.17$ | $0.58 \pm 0.18$ |
| Abstractness | High Arousal | $0.42 \pm 0.24$ | $0.45 \pm 0.20$ | $0.50 \pm 0.25$ |
| | Low Arousal | $0.13 \pm 0.19$ | $0.13 \pm 0.14$ | $0.21 \pm 0.21$ |
| *PC1:LenFreq* | High Arousal | $0.29 \pm 1.32$ | $0.66 \pm 0.91$ | $-0.13 \pm 0.91$ |
| | Low Arousal | $-0.41 \pm 0.83$ | $-0.05 \pm 0.79$ | $-0.36 \pm 0.69$ |
| *PC2:Valence* | High Arousal | $-0.22 \pm 1.27$ | $-0.36 \pm 1.48$ | $-0.54 \pm 0.80$ |
| | Low Arousal | $0.44 \pm 0.71$ | $0.35 \pm 0.48$ | $0.33 \pm 0.27$ |
| *PC3:Abstractness* | High Arousal | $0.33 \pm 1.02$ | $0.49 \pm 0.71$ | $0.68 \pm 1.04$ |
| | Low Arousal | $-0.65 \pm 0.83$ | $-0.58 \pm 0.69$ | $-0.27 \pm 0.81$ |

As is evident from the table, a perfect balance in the control variables was not possible to achieve. Specifically, High Arousal words tended to be somewhat longer, more negative, more abstract and less frequent than Low Arousal words, and this pattern appeared consistent across all three languages. We therefore decided to use principal components of the control variables as additional covariates in subsequent analyses. As explained in more detail in Appendix S2, the original control variables were condensed into three principal components, labelled *PC1:LenFreq, PC2:Valence*, and *PC3:Abstractness*, respectively; a summary of their Anderson-Rubin factor scores (Means and *SDs* across items) is provided at the bottom of Table 1. Using principal components in later analysis not only resulted in a more manageable number of control predictors, but also made their contributions to the model fit easier to estimate (principal components are uncorrelated to one another). Across the 180 items, a multiple regression analysis predicting *Word Type* (coded as 0=Low Arousal, 1=High Arousal) from the three principal components yielded *R^2^* = .47, which translates into a *Variance Inflation Factor* of 1.9 and thus tolerable levels of collinearity with the predictor variable of interest (according to the least liberal convention in the regression literature, *R^2^* > .6 / *VIF* > 2.5 would be regarded as potentially problematic).

For each language, we also selected 30 filler words with ‘medium’ normalised arousal scores (Mean ± *SD:* 0.39 ± 0.02) but from a generally lower lexical frequency range (0.59 ± 0.53) than the experimental items. Care was taken to ensure that the filler materials did not differ systematically in terms of length, frequency, valence, or abstractness across languages.

### Procedure

At the beginning of each experimental session, bilingual participants first filled in the LHQ questionnaire. Native English speakers were simply asked whether they used any foreign languages regularly; all of them confirmed that English was the only language they produced and comprehended on a day-to-day basis. After a brief eye-dominance test (41), participants were seated ca. 70cm from a 21-inch CRT monitor (100 Hz refresh rate, 1024 × 768 pixel resolution) connected to an SR-Research EyeLink II head-mounted eye-tracker (0.01° spatial resolution) running SR-Research ExperimentBuilder software. Viewing was binocular, but only the participant’s dominant eye was tracked, using a sampling rate of 250 Hz. After initial camera setup, the eye-tracker was calibrated using the standard EyeLink nine-point calibration routine. Setup and calibration usually took less than a minute to complete. Calibration was repeated at least 3 times during a recording session. English monolingual participants were tested in a single recording session each (typically lasting no more than 15 minutes) using English target materials only. For the bilingual participants, there were two separate recording sessions (interspersed by a 5-10 minute break), as they were tested both in their first language (Finnish respectively German) and in their second language (English). Ordering of recording sessions (L1 first or English first) was counterbalanced across participants, and each recording session started with a brief reminder of the instructions in the appropriate target language (English, Finish, or German, respectively).

Within each recording session, 90 word-stimuli (30 high arousal, 30 low arousal, and 30 fillers) appeared in an individually determined quasi-random order, subject to the constraint that the first four trials per session were fillers. As illustrated in Fig 1, each individual trial started with the presentation of a central fixation dot for drift correction, followed by a 11-character mask of ‘X’es (duration: 500 ms), the actual word (duration: 50 ms + 20 ms × length), and then the mask again (duration: 1700 ms). Note that the presentation duration of the word was a function of its length (in characters) and ranged from 110 ms to 250 ms across trials. At the end of each trial, a blue question mark prompted participants to provide a word recognition judgement, i.e. to press the right-hand trigger (for “*yes, I recognised the word and its meaning*”) or the left-hand trigger (for “*no, I did not recognise the word or its meaning*”) on a hand-held Microsoft USB game pad used as a button box throughout the experiment. After providing the button response, the next trial was initiated after a random delay of 200-500 ms. The words, as well as the mask of ‘X’es, appeared centrally on screen in 28-point Monospace capital letters, printed in black on a light grey background (RGB 230,230,230). Participants were instructed to keep looking at the centre of the screen throughout each trial, and to perform eye-blinks ‘in between’ trials if possible.

**Fig 1.**
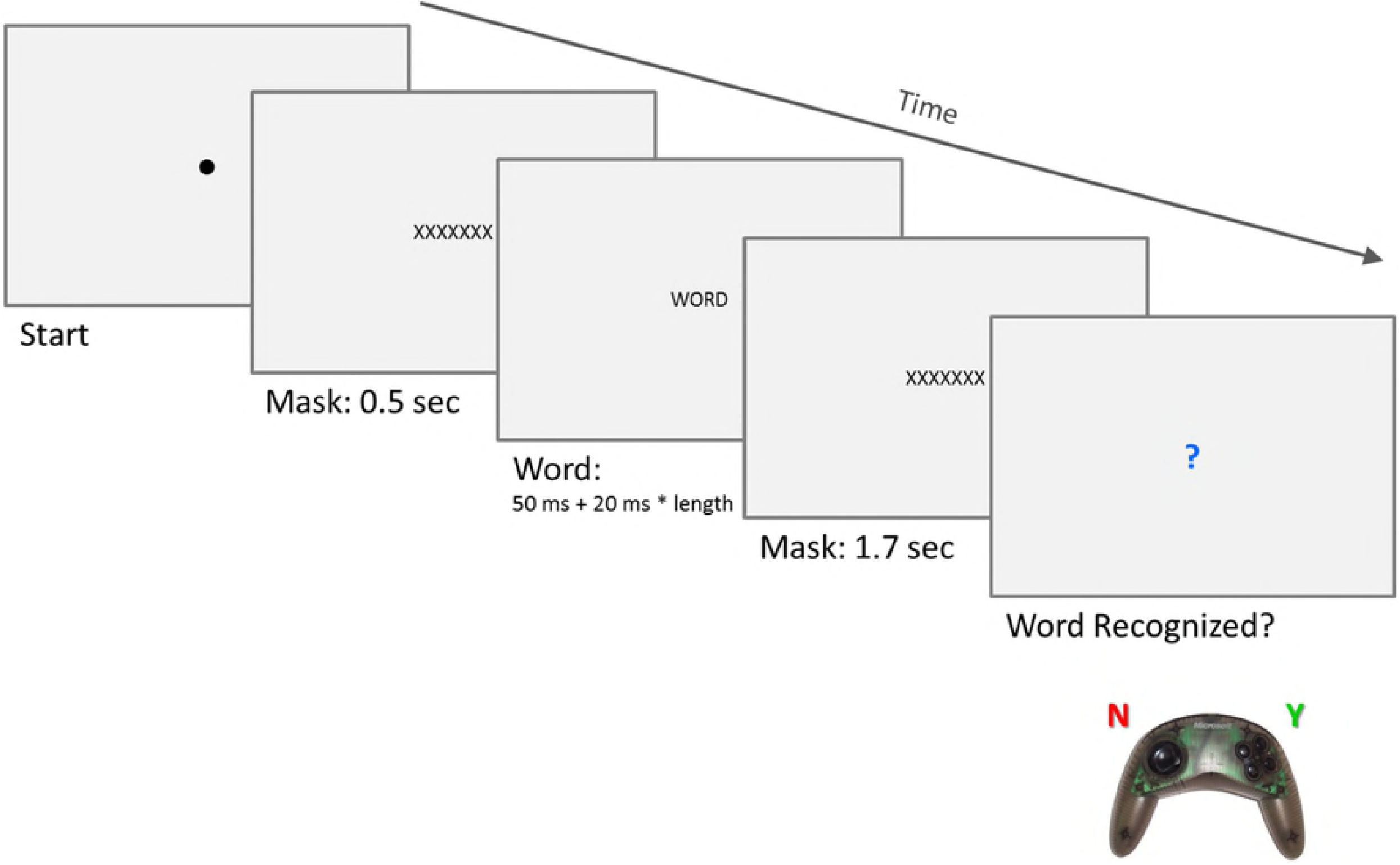
Presentation sequence per trial. The recording sessions took place in an experimental room with blacked-out windows. Surrounding luminance conditions were kept constant using fluorescent ceiling lights.

## Analysis and Results

### Off-line Word Recognition Judgements

Word recognition judgements at the end of each trial were scored as either 0 (word not recognized) or 1 (word recognised). These data were aggregated into mean probabilities of affirmative judgements, broken down by participant, target language, and word type. The mean probabilities were then subjected to non-parametric bootstrapping over 10,000 resamples (repeated-measures data from the same participant were treated as one unit for resampling) to determine by-subject 95% confidence intervals in each *Target Language* × *L1* × *Word Type* combination. The results are shown in Fig 2.

**Fig 2.**
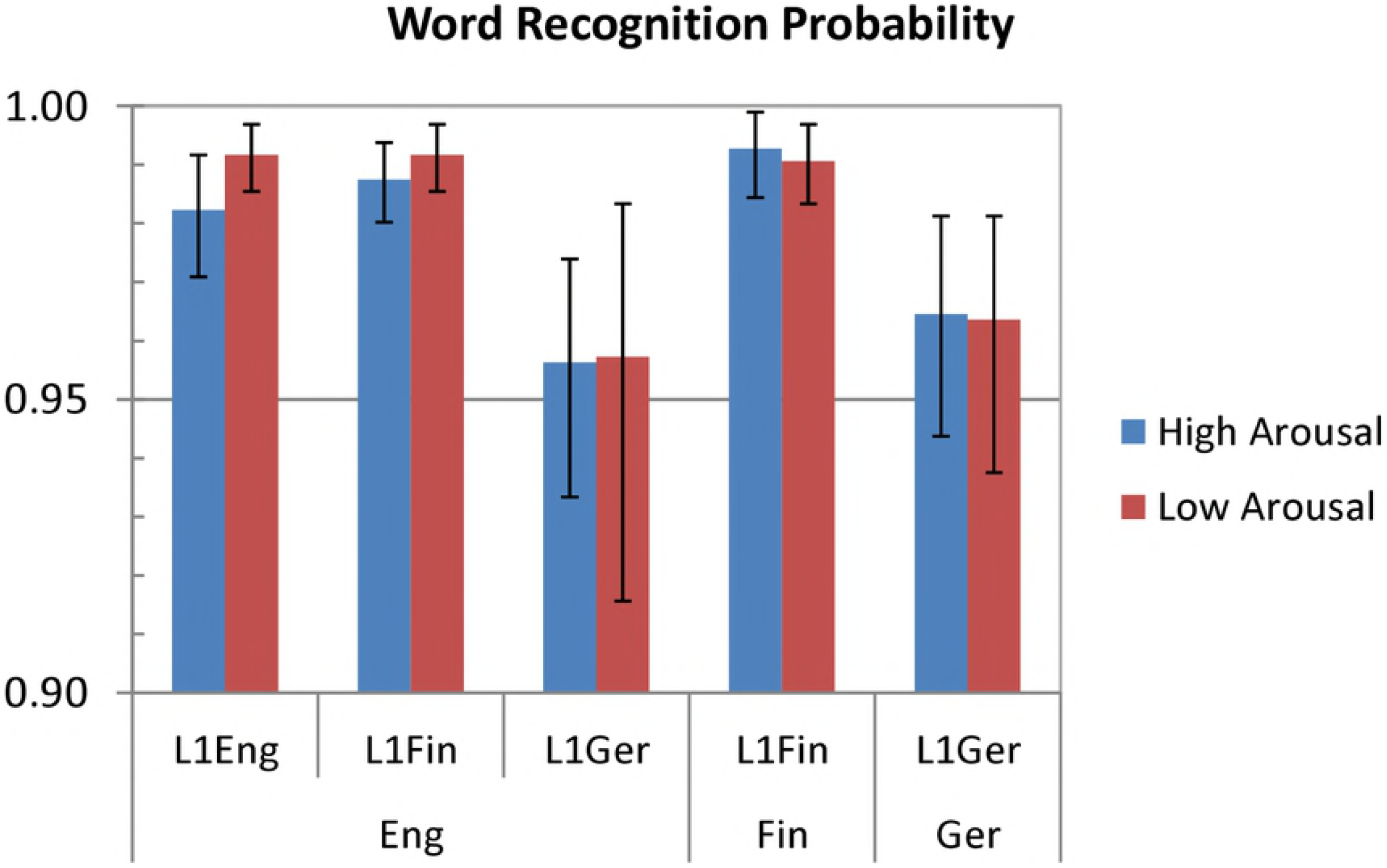
Probabilities of affirmative word recognition judgements. Broken down by Target Language (English, Finnish, or German – bottom labels on the X-axis), L1 of Participant (English, Finnish, or German), and Word Type (high arousal vs. low arousal). Error bars indicate bootstrapped 95% confidence intervals by participants.

Fig 2 indicates that the likelihood of an affirmative word recognition judgement was well above 90% in each design cell. There were no appreciable differences related to *Word Type*. Compared to the English and Finnish participants, native German bilinguals were generally more conservative (and more variable) in providing an affirmative response, not only when tested in their L2 (English) but also in their L1 (German). The native Finnish bilinguals were virtually indistinguishable from the English monolinguals when tested in English, and also when tested in Finnish. Overall, these data suggest that participants could easily recognise the stimuli in each *Word Type* condition, regardless of whether they were tested in their first or in their second language.

## Pupil Size Data

### Data Pre-processing

The eye-tracker continuously recorded participants’ pupil-sizes and eye-positions over the entire duration of each trial, using a sampling rate of 250Hz. For each participant and trial, we extracted pupil size and eye-position data for a time period starting from 100 ms before the onset of the critical word presentation and ending at 1900 ms after the onset of the critical word presentation. To make the data more manageable in size, this 2-second time period was broken down into twenty time bins of 100 ms each (down-sampling to 10Hz resolution).

For each of the twenty critical time bins per trial, we extracted the average pupil size (area in pixels per eye-camera sample) and eye-position on screen (average X- and Y-coordinates in pixels) from the eye-tracker output, using SR-Research Data Viewer software (Version 2.1.1). Missing data due to eye-blinks (less than 1.7% of all data points, randomly distributed across participants, trials and time bins) were replaced with linearly interpolated estimates from temporally adjacent time bins; a total of 12 trials (from 4 different participants) were removed altogether because they contained blink-related gaps of more than half a second in duration. The pupil size data, but not the eye-position data, were converted into decimal logarithms for further analysis; differences on the log scale correspond to proportional changes on the raw scale. Although participants were instructed to fixate the centre of the screen throughout the entire trial period, small eye-movements or drifts (rarely exceeding 2 degrees of visual angle) were not uncommon, which induced a certain amount of noise in the pupil size data. (Camera angles relative to the recorded eye varied across participants and recording sessions). To remove this kind of noise, we performed regression analyses with X- and Y-position of the eye as orthogonal predictors of log pupil size. This was done separately for each participant and recording session, and using all the data available (20 [time bins] × 90 [experimental and filler trials] = 1800 data points per participant and recording session). The predicted log pupil sizes from these regression analyses were then subtracted from the actual log pupil sizes per trial and time bin, i.e. subsequent analyses were based on position-adjusted *residual* log pupil size data.

### Simple Effects Over Time

In all analyses that follow, filler trials were excluded. Since reported word recognition rates were high (see Fig 2), we did not remove any trials based on off-line judgements. The position-adjusted log pupil size data per trial (cf. previous section) were converted into deviations from the *baseline* established within the time bin from 100 to 0 ms before the onset of the critical word presentation (this baseline time bin is labelled “−100” in the plots that follow). For each of the twenty 100-ms time bins per trial, we thus ended up with a score, labelled Δ*BL*, that quantified the difference between the current vs. the *baseline* log pupil size (both adjusted for eye-position). By then calculating 10^Δ*BL*^, we obtained a measure of the *proportional* change in pupil size relative to the baseline, such that a score of (say) 1.05 would indicate a 5% increase and a score of 0.95 a 5% decrease in pupil size relative to the baseline; at time bin “−100” (the actual baseline time bin), this score was always equal to 10^0^ = 1.

In a first, more descriptive analytical step, we explored the simple effect of *Word Type* on these proportional changes from baseline pupil size, separately for each participant sample and recording session: (1) English monolinguals tested in English; (2) Finnish bilinguals tested in Finnish; (3) Finnish bilinguals tested in English; (4) German bilinguals tested in German; and (5) German bilinguals tested in English. The analyses were based on linear mixed effects models (42) and were further split by time bin to examine pupil size changes over time. (The latter is not ideal to account for autoregressive dependencies in time series data, but should nevertheless be descriptively informative.)

The linear mixed model (LMM) analyses per time bin were carried out in SPSS (v21) and included *Word Type* (High vs. Low Arousal) as the only fixed effect, together with various crossed random effects to simultaneously account for by-subject and by-item variability in the data. Following (43), the maximal random effect structure justified by the design was used. That is, we included *Subject* and *Item* as random main effect terms (or *random intercepts* in LMM terminology) and *Subject × Word Type* (the *by-subject random slope on Word Type*) as the only random interaction term. This appropriately models observational dependencies resulting from the within-subjects/between-items nature of the *Word Type* manipulation. The only deviation from the maximal approach was that the models did not include any random correlation terms (models with random correlations led to convergence problems). Finally, we also included the three principal components derived from the item-related control variables (*PC1:LenFreq, PC2:Valence*, and *PC3:Abstractness*) as additional random covariates in the analysis models. The purpose of this was to adjust estimated marginal means and standard errors for these control predictors, i.e. to establish the simple effect of *Word Type* after accounting for potential effects of the control variables on pupil size. We included the control predictors as random effects because their value distributions were dependent on the actual items used (if the study were replicated with new sets of words per language, these value distributions would be different). We also performed analyses without the covariates, which resulted in generally stronger effects of *Word Type* on pupil size. The latter suggests that some variability in pupil size could indeed be attributed to moderate cross-condition differences in length, frequency, valence, and abstractness, which were not of primary theoretical interest.

The LMM parameter estimates were used to determine standard errors for the difference in pupil size between high- versus low-arousal words (i.e., the simple effect of *Word Type* after accounting for subject, item, and control predictor variability). Fig 3 shows the results, separately for each participant group and recording session: Across conditions, pupil size rose to a maximum of ca. 3% above baseline at around 1 second after stimulus onset, before dropping again (to just under baseline level) towards the end of the considered time period. More critically, Fig 3 indicates that words associated with low emotional arousal engendered significantly less dilated pupils than words associated with high emotional arousal, but only when participants were tested in their first language (panels on the left) and not when participants were tested in their second language (panels on the right). In the latter case, simple effects of *Word Type* on pupil size did not reach significance in any of the time bins considered. For English monolinguals, the *Word Type* effect emerged from time bin “700” (milliseconds post stimulus-onset) onwards; Finnish bilinguals (tested in Finnish) showed the effect from time bin “600” onwards, and German bilinguals (tested in German) from time bin “900” onwards.

**Fig 3.**
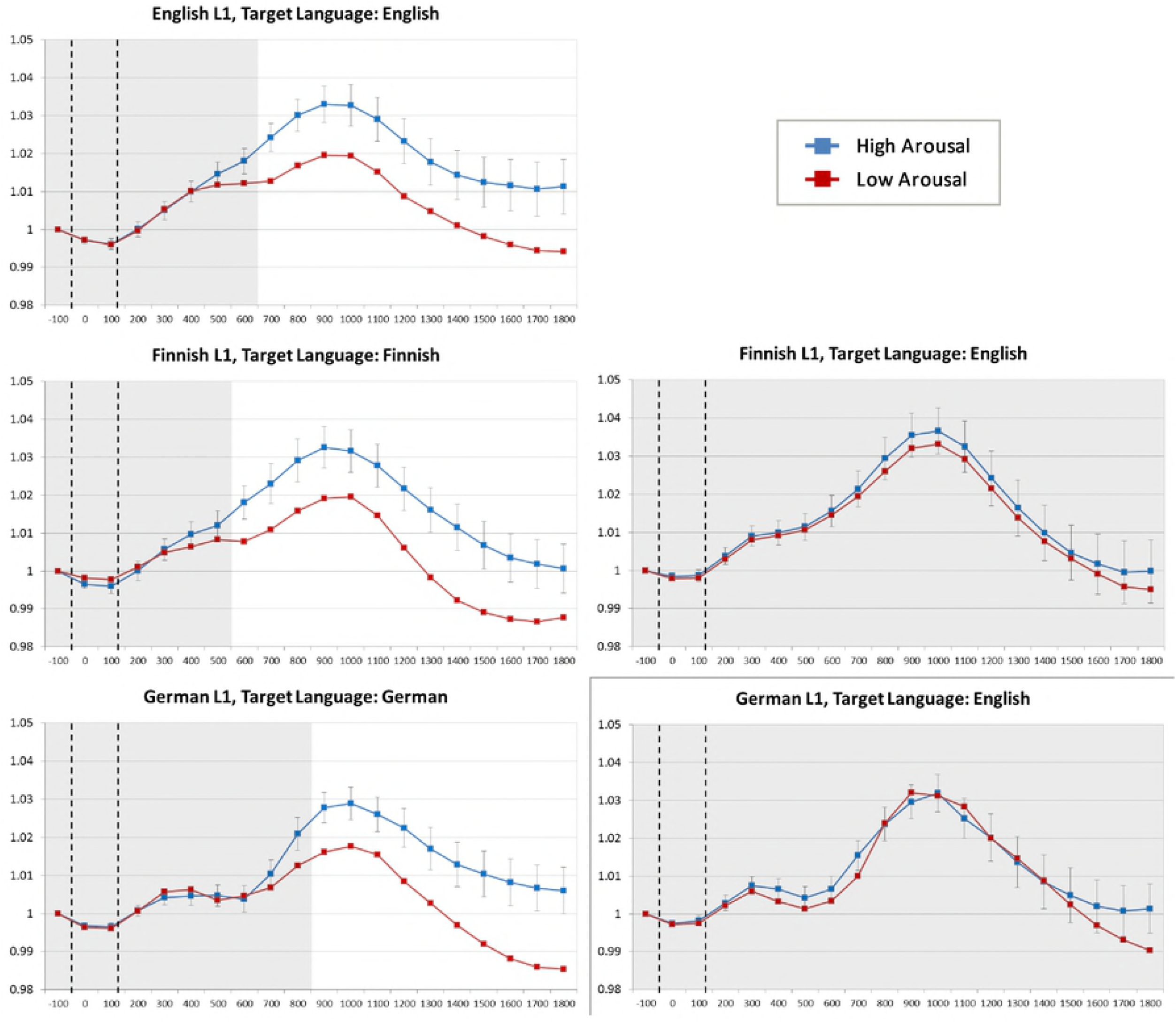
Covariate-adjusted estimated marginal means for proportional pupil size changes over time. Time is partitioned into 100-ms time bins (X-axis labels refer to the temporal onset of each time bin relative to stimulus onset). Vertical dashed lines indicate the average onset and offset of the presented word. Blue curves: High Arousal words; red curves: Low Arousal words; Top to bottom: English, Finnish, and German L1 participants; Left hand panels: stimuli presented in participants’ first language; Right hand panels: stimuli presented in participants’ second language. Error bars refer to standard errors for the simple effect of Word Type derived from linear mixed effect model analyses (see text). White areas indicate time bins where the simple effect of Word Type was significant at *p* < .05.

### Area-Under-Curve Analyses

To inferentially test whether and how effects of *Word Type* on pupil size interacted with participants’ L1 and/or the language they were tested in, the data per trial were condensed into sum-scores to allow for area-under-the-curve comparisons. Specifically, we added pupil size data from time bin “600” onwards together, and multiplied the result with 100, thereby obtaining an estimate of the area under the curve (in millisecond × proportional pupil-change units) for the time period from 600 ms to 1900 ms after stimulus onset. This was done for each individual trial, resulting in one area under the curve score for each subject × item combination. Apart from time bin “600” being the earliest where a simple effect of *Word Type* emerged (see previous section), the motivation behind choosing this time bin as area starting point was the following general observation (see Fig 3): within the first 0-600 ms from word onset, pupil-size first dropped to just under baseline level, and then rose again to just above baseline level, reaching a temporary plateau by around 500-600 ms from stimulus onset (time bin “500”). We conjecture that this initial pattern could reflect a pupillary response to the changing stimulus (*mask* > *word* > *mask*) which is unrelated to word processing per se.

The area-under-the-curve data were then entered into two different types of LMM analyses. The first focused on bilingual participants only and examined potential interactions between the within-subjects effect of *Word Type* (High vs. Low Arousal), the between-subjects effect of *L1* (Finnish vs. German), and the within-subjects effect of *Target Language* (L1 vs. English). The second type of LMM analysis considered English target language data only, and compared effects of *Word Type* on pupil size across the three groups of participants (English monolinguals, Finnish bilinguals, and German bilinguals).

For the bilinguals-only analysis, the area under the curve data were entered into a LMM with a full-factorial *L1* (Finnish vs. German) × *Target Language* (L1 vs. L2) × *Word Type* (High vs. Low Arousal) fixed effects design. Additionally, the model included *Subject* and *Item* as random main effect terms, the *Subject × Word Type, Subject × Target Language*, and *Subject × Target Language × Word Type* random interactions (to account for by-subject variability in effects involving *Target Language* and *Word Type*, both of which were within-subjects), the three random covariates *PC1:LenFreq, PC2:Valence* and *PC3:Abstractness*, and finally, the interaction between *Target Language* and each of the three covariates as separate 2-way random interaction terms (to account for potentially different slopes of the covariates dependent on whether participants were tested in their L1 or in their L2). All random effects involving *Subject* were nested within *L1*, and the *Item* random main effect was nested within *Target Language*. Table 2 shows the fixed effects results, and the corresponding random effect estimates are provided in Appendix S3.

**Table 2.**
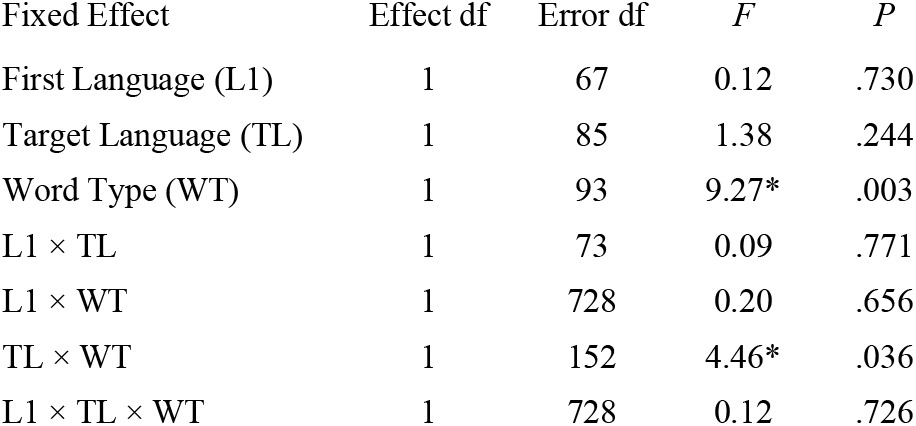
Fixed effects results from LMMs considering bilingual participants only. Area-under the curve data were modelled by fixed factor combinations of participants’ L1 (Finnish vs. German), Target Language (L1 vs. L2), and Word Type (High vs. Low Arousal). Significant *F*-values (Type III variance decomposition) are marked with an asterisk.

| Fixed Effect | Effect df | Error df | *F* | *P* |
| --- | --- | --- | --- | --- |
| First Language (L1) | 1 | 67 | 0.12 | .730 |
| Target Language (TL) | 1 | 85 | 1.38 | .244 |
| Word Type (WT) | 1 | 93 | 9.27* | .003 |
| L1 × TL | 1 | 73 | 0.09 | .771 |
| L1 × WT | 1 | 728 | 0.20 | .656 |
| TL × WT | 1 | 152 | 4.46* | .036 |
| L1 × TL × WT | 1 | 728 | 0.12 | .726 |

The fixed effects results (Table 2) confirmed a significant *Word Type* main effect, which was further modulated by a reliable *Target Language* × *Word Type* interaction. The corresponding estimated marginal means and 95% confidence intervals (Fig 4) indicate that the Effect of *Word Type* on pupil size was reliable only when bilingual participants were tested in their L1 (Finnish or German), but not when they were tested in their L2 (English). Since the 3-way interaction did not approach significance, this pattern did not depend on whether participants were native Finnish or native German speakers.

**Fig 4.**
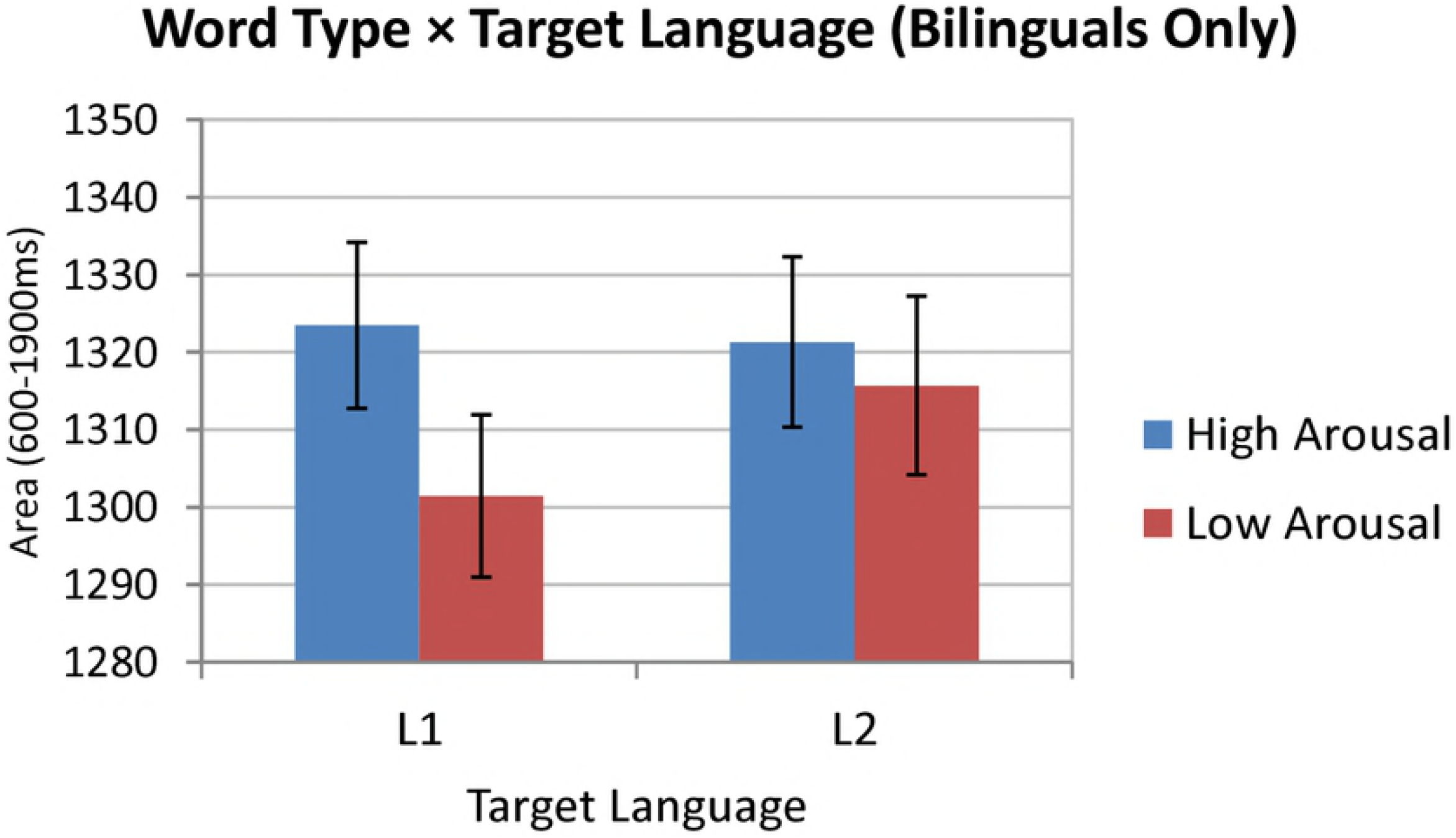
Estimated marginal means (with 95% CIs) for the Target Language × Word Type interaction. (Analysis considering bilingual participants only). Dependent variable was the pupil size area under the curve from 600-1900ms after stimulus onset. Means and CIs are covariate-adjusted.

The second LMM analysis focused on English target language data only. The area-under-the-curve scores were entered into a model with a full-factorial *L1* (English vs. Finnish vs. German) × *Word Type* (High vs. Low Arousal) fixed effects design. As random effects, we included the main effect of *Subject* (nested within L1), the main effect of *Item*, the *Subject* × *Word Type* random interaction (nested within L1), the three random covariates (*PC1:LenFreq, PC2:Valence*, and *PC3:Abstractness*) and their 2-way interactions with *L1* (the latter accounting for potentially different slopes of the covariates dependent on participants’ first language). Table 3 summarises the fixed effects results (see Appendix S3 for the corresponding random effects estimates).

**Table 3.**
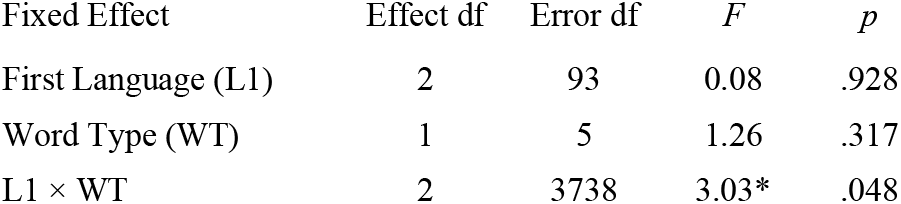
Fixed effects results from LMMs considering English target language only. Area-under the curve data were modelled by fixed factor combinations of participants’ L1 (English vs. Finnish vs. German) and Word Type (High vs. Low Arousal). Significant F-values (Type III variance decomposition) are marked with an asterisk.

As Table 3 shows, the only significant fixed effect (at just under .05 level) was the *L1 × Word Type* interaction. The corresponding estimated marginal means and 95% CIs are shown in Fig 5, indicating that when participants were tested in English, the effect of Word Type on pupil size was detectable only in English monolinguals, but not in Finnish or German bilinguals.

**Fig 5.**
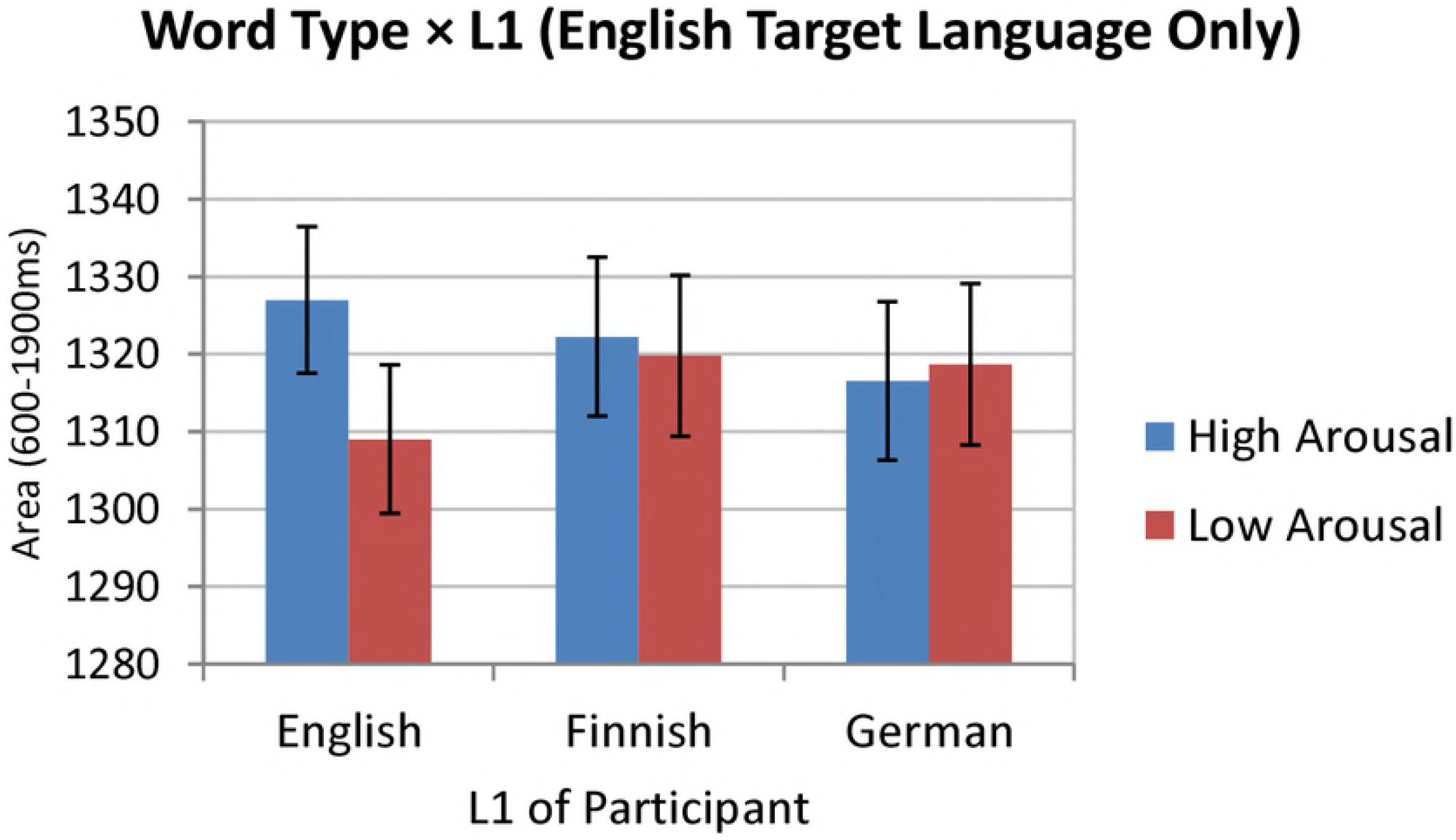
Estimated marginal means (with 95% CIs) for the L1 × Word Type interaction. (Analysis considering English target language only). Dependent variable was the pupil size area under the curve from 600-1900ms after stimulus onset. Means and CIs are covariate-adjusted.

## Discussion

The aim of the present pupillometry study was to further corroborate and theoretically extend previous findings on reduced emotional resonance in L2. Specifically, it was hypothesised that differences in pupillary responses to high-versus low-arousing words should be more pronounced in bilinguals’ L1 (here, German or Finnish) than in their L2 (here, English). This hypothesis was clearly supported by our data: for the German and Finnish bilingual groups, a significant effect of word type on pupillary responses was found only when the stimuli were presented in their respective L1. When the same participants were tested in L2 (English) there was no reliable effect of word type on pupil size, resulting in a significant *Word Type × Target Language* interaction in the *bilinguals only* analysis of pupil size (cf. Fig 4). Importantly, comparisons with a monolingual English control group of participants suggested that the English stimuli were no less effective than the German or Finnish stimuli in eliciting affective pupillary responses. Specifically, the *English target language only* analysis registered a reliable *L1 × Word Type* interaction whereby high-arousing words in English evoked more dilated pupils than low-arousing words, but only in L1 speakers and not in bilingual L2 speakers of English (cf. Fig 5). Taken together, the present pupillometry data in support of reduced emotional resonance in L2 can be regarded as an important extension and confirmation of previous demonstrations where target language varied *between-subjects* and where no monolingual control group was included (cf. (32)).

A further advancement, compared to previous studies, is that we can be reasonably confident in ruling out word recognition difficulties in L2 as a potential confound: Participants provided word recognition judgements after every individual trial per experimental session, and there was no indication that our bilingual participants (all of whom reported to be highly proficient in English) were struggling to process materials presented in L2. (Importantly, while German participants tended to be more conservative in their judgements, this was independent of the language they were tested in, see Fig 2). Combined with the fact that potential word-related confounds such as length, frequency, and abstractness were controlled for (both analytically and by design) this leaves reduced emotional resonance as the most likely explanation of the observed reduction in pupillary response contrasts between high- and low-arousing words in L2.

A point worth noting is that our study did not find L2 word processing to be associated with generally more dilated pupils than L1 word processing, as suggested by a non-significant *Target Language* main effect in the *bilinguals only* analysis (Table 2). While in line with the L1 vs. L2 comparisons in (32) (which were made *between-subjects*), the lack of a clear *Target Language* main effect in our study does not support the hypothesis that processing of L2 would pose generally higher cognitive demands than processing of L1. On the other hand, Fig 3 and Fig 4 do show clear increases in pupil size for low-arousing words in L2 relative to L1, whereas pupil sizes for high-arousing words remained largely comparable across *Target Language* conditions. Unfortunately, due to the lack of a suitable emotional baseline condition – a potential shortcoming that our investigation shares with previous studies – it is difficult to assess the exact contribution of cognitive load versus emotional resonance in determining pupillary responses to materials in L2. (Note that the filler words in our study do not constitute an ideal baseline because, among other things, they were from a generally lower lexical frequency range than the critical items). Given the observed pattern, however, it would seem premature to rule out increased cognitive load as a potential contributor to pupillary responses in L2 ‘on top of’ (or perhaps even interacting with) affective influences on pupil size. This remains an important topic for future research, which would ideally include an affectively ‘neutral’ baseline condition.

Another noteworthy aspect of the present findings concerns linguistic distance between L1 and L2. The present did not provide any compelling evidence in support of a modulating influence of this factor on reduced emotional resonance in L2. Specifically, when L1 and L2 were more closely related in comparative linguistics terms (i.e., German vs. English), the L2-related reduction in the pupil size contrast between high- and low-arousing words was more or less the same as when L1 and L2 were more distant (i.e., Finnish vs. English) – see Fig 3 and the non-significant 3-way interaction in Table 2. This may be taken to suggest that reduced emotional resonance in L2 is not dependent on similarities in linguistic features (e.g., basic vocabulary, phonology, and morphology), but is rather a question of linguistic experience (later, more ‘formal’ acquisition of L2 or less exposure to L2 over one’s lifetime).

Overall, our results fall in line with previous findings based on skin-conductance (e.g., (20–22)) as well as pupillometry (32), suggesting that emotion word processing, and specifically emotional resonance in L1 vs. L2, is linked to activation in the Autonomic Nervous System. Thus, our study contributes to a growing body of work suggesting that (bilingual) word processing is partially grounded in physiological reactions to emotion, and that reduced emotional resonance can be detected with physiological measures.

One concern might be that we did not explicitly control for L2 proficiency in our sample of bilinguals, which is likely to affect cognitive effort in processing L2. However, all of our bilingual participants (a) self-reported to be highly proficient in English, (b) were Glasgow University students using English as their language of study, and (c) had been living in an English-speaking country for minimum of three months (average: two years and four months) when the study was conducted. Together with the fact that post-trial word recognition rates were well above 90% across participant groups and conditions (Fig 2), we do not believe that L2 proficiency had a major impact on our results. That said, we agree that proficiency should ideally be controlled for in future research, e.g. by adding a suitable measure as another person-specific covariate in the analyses. Previous research had identified proficiency as a potential mediating factor of perceived emotionality in the Bilingualism and Emotions Questionnaire (44), and it had also been suggested that bilinguals with close-to-native proficiency in L2 show similar affective processing in L1 and L2 (16).

In relation to pupil-size changes, one aspect that our current data are unable to address is whether reduced emotional resonance primarily affects the *strength* (amplitude) or the *timing* of pupillary responses to high- vs. low-arousing words in L2 (note that in (11), for instance, pupil-size changes in response to varying degrees of word-retrievability were delayed in L2 compared to L1). Indeed, our area-under-the-curve analyses are not very informative in this respect, but a closer inspection of the plots in Fig 2 suggests that pupillary responses generally peaked around 900–1000 ms after word onset and, more interestingly, that temporal peak locations remained remarkably consistent across *L1 × Target Language × Word Type* conditions. We therefore lean towards concluding that the affective effects registered in our study were largely driven by differences in *amplitude* rather than timing of pupillary responses. On the other hand, particularly for the German bilinguals tested in English, there was a suggestion that the two *Word Type* conditions started to separate – in the expected direction – towards the end of the considered time period per trial (bottom-right panel of Fig 2), which may be taken to indicate that the *Word Type* effect was considerably delayed in L2 relative to L1. Unfortunately, without measuring pupillary responses over longer periods of time after stimulus-onset (which would arguably be more taxing to participants), it appears difficult to obtain a definitive answer to this interesting question. At the very least, our data confirm a change in pupillary responses to high- vs. low-arousing words in L2 relative to L1. This change may be characterised in terms of smaller differences in response amplitudes (i.e., ‘weaker’ affective effects) or in terms of ‘delayed’ affective effects in L2. While we cannot conclusively rule out the latter possibility, we think that the former interpretation is more plausible given the observed consistency in the timing of peak responses.

To conclude, the present study evaluated pupillometry as a relatively novel physiological method of establishing emotional resonance in late bilinguals’ first versus second language. In line with previous findings (e.g.,(19–21, 32)), our data show a less pronounced contrast in affective physiological responses to high- vs. low-arousing words when materials were presented in participants’ second rather than first language. Importantly, influences of eye-movements and of potential lexical confound variables were carefully controlled for in our study, and there was no indication that bilinguals were struggling to identify the words in L2. Apart from strengthening the evidence for reduced emotional resonance in L2, our study further suggests that linguistic relatedness between L1 and L2 plays no major role in determining affective response patterns across languages. Overall, the present findings, and those by (32), indicate that pupillometry is a promising alternative to skin-conductance research when measuring direct physiological responses to emotional content in different languages. Future research may use a combination of these methods to answer theoretical questions related to the causes of reduced emotional resonance in L2.

